# Climatic and non-climatic drivers of rangeland vegetation change in Nepal

**DOI:** 10.64898/2026.07.09.737421

**Authors:** Uttam Babu Shrestha, Suraj Joshi

## Abstract

Nepal’s rangelands provide multiple benefits, including support for pastoral livelihoods and alpine biodiversity, regulation of water and soil nutrients, and sequestering carbon. Climate change and anthropogenic pressures are altering these rangelands, leading to vegetation and biodiversity change. However, national-scale assessments of rangeland change are limited in Nepal. This study quantified rangeland changes at multiple spatial scales and assessed the climatic and non-climatic drivers of rangeland change. About 80.7% of Nepal’s high-altitude rangeland (> 2,000m) outside protected areas showed no significant change. Among areas exhibiting significant annual maximum NDVI trends, 383,281 ha (18.6%) showed positive and 14,702 ha (0.7%) showed negative trends, corresponding the ratio of increase in vegetation greenness and decline in vegetation greenness to 26:1. Climate predicted positive trends covered 627,184 ha (30.5%), whereas residual trends caused by non-climatic drivers covered 94,656 ha (4.6%). Climate induced negative trends covered 47,609 ha (2.3%) while residual trends were observed in 6,260 ha (0.3%). Negative trend pixels were concentrated mainly within the 3,000–5,000 m elevation band, with Karnali Province recording the highest proportional climate predicted decline in vegetation greenness (3.4%). At the municipality scale, rangeland change showed no significant relationship with grazing pressure derived from gridded livestock data, suggesting that grazing pressure alone did not explain the non-climatic vegetation signal. These spatially explicit, nationally consistent results identify where rangeland change is occurring and help distinguish climatic and non-climatic drivers of rangeland vegetation change, providing evidence to support targeted rangeland management under Nepal’s federal governance structure.

## 1. Introduction

Rangelands, encompassing grasslands, savannas, shrublands, steppes, deserts, and alpine meadows, are among the most expansive terrestrial biomes on Earth, covering approximately 54% of the global land surface and supporting critical ecological functions and human livelihoods (ILRI et al. 2021; Reeves et al. 2024; Xie et al. 2024). Rangelands are often characterized by low and erratic rainfall, rugged terrain, poor drainage, and low temperatures, which make them unsuitable for cultivation (Getabalew & Alemneh 2019). These ecosystems are primarily utilized for livestock grazing by hundreds of millions of pastoral and agro-pastoral people worldwide, supporting food security, income, and cultural identify (Reed et al. 2015; Seid et al. 2016; Godde et al. 2020; Dean et al. 2021). Beyond livestock production, rangelands contribute to nutrient cycling, water regulation, carbon sequestration and storage, maintenance of biodiversity including endemic and threatned species, and the preservation of cultural heritage and spiritual values embdedded in pastoral communities (Seid et al. 2016; Dean et al. 2021; Maczko et al. 2022; Bennett et al. 2023). Despite their ecological and socio-economic importance, rangelands are increasingly threatned by climatic and non-climatic drivers, which negatively affect ecosystem health and the resilience of pastoral communities (Maczko et al. 2022; Xie et al. 2024; Simba et al. 2024).

Rangeland degradation is one of the most pressing and widespread environmental challenges globally although its definition is contested and context-dependent across ecological and governance frameworks (Dregne 2002; Bedunah & Angerer 2012; Silcock & Fensham 2019; Sainnemekh et al. 2022). Rangeland degradation generally refers to a sustained decline in the productive capacity and ecological functioning of rangeland systems, often reflected in reduced vegetation cover, forage biomass, soil organic carbon, and plant species diversity (Pickup et al. 1993; Milton et al. 1994). Major documented drivers of rangeland degradation include overgrazing and livestock intensification, land-use conversion through cropland expansion, urbanization, infrastructure development, and large-scale tree planting in grassy systems, woody plant encroachment, invasive species, climate-induced drought, and shifts in precipitation regimes (Seid et al. 2016; Burrell et al. 2019; Bardgett et al. 2021; Sainnemekh et al. 2022; Xie et al. 2024; Briske et al. 2024). Coupled with these drivers, pastoral abandonment and the weakening of traditional governance systems have further reduced the adaptive capacity of both rangeland ecosystems and pastoral communities (Bedunah & Angerer 2012; Sainnemekh et al. 2022).

In Nepal, rangelands occupy approximately 14.7% - 22.6% of the total land area (FRTC 2022; GoN 2012), with more than 90% concentrated in the mountain and Himalayan zones above 2,500 m elevation, where they provide the primary source of fodder for high-altitude livestock (LRMP 1986; Pande 2009). These high-altitude grazing systems also support significant biodiversity, including endemic alpine flora, economically valuable medicinal and aromatic plants, and habitat for globally significant wildlife (Dong et al. 2009; MoITEF 2026). In addition to supporting mountain livelihoods and biodiversity, Nepal’s rangelands play a crucial role in providing watershed services, carbon sequestration, supporting ecotourism, and promoting cultural identity, particularly through Himalayan transhumance systems (Aryal et al. 2014; MoITEF 2026).

However, Nepal’s rangelands face multiple intersecting pressures. In many areas, livestock stocking rates are substantially higher than estimated carrying capacities, leading to overgrazing (Barsila 2008; Pande 2009; Parajuli et al. 2013). Field-based studies show that grazing intensity profoundly influences vegetation structure, plant species diversity, biomass production, and soil physicochemical properties in Nepal’s rangelands although the direction and magnitude of these effects vary substantially by elevation and site-specific conditions (Chaudhary et al. 2025). Socio-economic changes including rural out-migration, shifting cultural values, and declining youth interest in herding, have contributed to pastoral abandonment in some areas, promoting shrub encroachment and the conversion of productive semi-natural grasslands into shrubland (Sharma et al. 2014; Gentle & Thwaites 2016; Tiwari et al. 2020). Additionally, climate change adds further pressure as Nepal’s high-elevation regions, where rangelands are most extensive, are warming faster than lowland areas (Shrestha et al. 2019; MoITEF 2026). Satellite-based phenological analyses have reported delayed start of the season and lengthening of the growing season in alpine and subalpine vegetation in Nepal, with moisture playing a dominant role in phenology and growing-season dynamics (Katel et al. 2026). Nevertheless, the combined effects of climatic and non-climatic drivers of Nepal’s rangelands remain poorly understood.

Despite growing recognition of the ecological and socio-economic pressures affecting Nepal’s rangelands, spatially explicit, long-term assessments that systematically distinguish climate-driven rangeland change from non-climatic residual change remain absent in Nepal except a few localized studies based on short-term remote sensing data (Paudel & Andersen 2010), field-based experiment (Chaudhary et al. 2025), and observational study (Banjade & Paudel 2008). Remote sensing offers a cost-effective, larger scale, and temporally consistent data for addressing this gap (Bai et al. 2008; Hill et al. 2008; Reeves et al. 2024). Satellite-derived vegetation indices, particularly the Normalized Difference Vegetation Index (NDVI) derived from the Moderate Resolution Imaging Spectroradiometer (MODIS) provide spatially continuous, multi-decadal records of vegetation productivity across ecosystems including remote and topographically complex regions such as the Himalaya (Eckert et al. 2015; Sanz et al. 2021).

Residual trend method (RESTREND) advances beyond simple NDVI trend analysis by regressing NDVI against climatic variable particularly precipitation to estimate the climate-predicted component of vegetation change (Burrell et al. 2017). The residual component is then used to identify vegetation trends not explained by the modeled climatic variables, such as land-use change, grazing pressure, shrub encroachment, management interventions and other non-climatic processes (Evans & Geerken 2004). In cold, high-altitude drylands as in Nepal Himalaya, including temperature alongside precipitation can improve the explanatory power of RESTREND and reduce the likelihood of incorrectly attributing vegetation change to non-climatic drivers (Burrell et al. 2019).

This study uses MODIS NDVI time-series data, ERA5 climate data, and RESTREND to assess rangeland vegetation dynamics across Nepal. Specifically, this study aims to a) characterize the spatial and temporal patterns of rangeland change across Nepal over a 26-year period; b) identify areas of significant increase (improvement) and decline (degradation) in vegetation greenness and their climatic and non-climatic drivers. The findings provide a spatially explicit evidence base to support targeted rangeland management, policy development, and conservation planning.

## 2. Materials and Methods

### 2.1. Study area

This study covered high elevation rangelands across Nepal, which lies between 26°22′N and 30°27′N latitude and 80°04′E and 88°12′E longitude. Nepal shares its northern border with the Tibetan Plateau and its southern border with Indo-Gangetic Plain of India. The country covers an area of 147,181 km² and spans one of the steepest elevation gradients on Earth, ranging from approximately 60 m above sea level (masl) in the Tarai lowlands to 8,849 m at the summit of Mount Everest. This elevation gradient creates diverse bioclimatic conditions, from tropical forests to alpine grasslands, and highly varied topographic and physiographic settings.

Rangeland extent was delineated using a moderate-resolution (30 m) annual land use and land cover (LULC) dataset for Nepal spanning 2000–2022 (FRTC 2022) (Figure 1). Rangeland pixels were extracted from each of the 22 annual LULC raster layers, and the annual classifications were combined to produce a temporally stable rangeland mask. Pixels classified as rangeland in more than six of the 22 years, representing at least 25th-percentile temporal consistency, were retained as stable rangeland pixels. This threshold was applied to minimize the inclusion of transitional or misclassified pixels while retaining genuine long-term rangeland extent. Since most rangelands in Nepal occur at higher elevations, the stable rangeland mask was further constrained to pixels above 2,000 m elevation using the Shuttle Radar Topography Mission (SRTM) 30 m digital elevation model (DEM; Jarvis et al. 2008). This final rangeland mask defined the spatial extent for all subsequent analyses.

**Figure 1.**
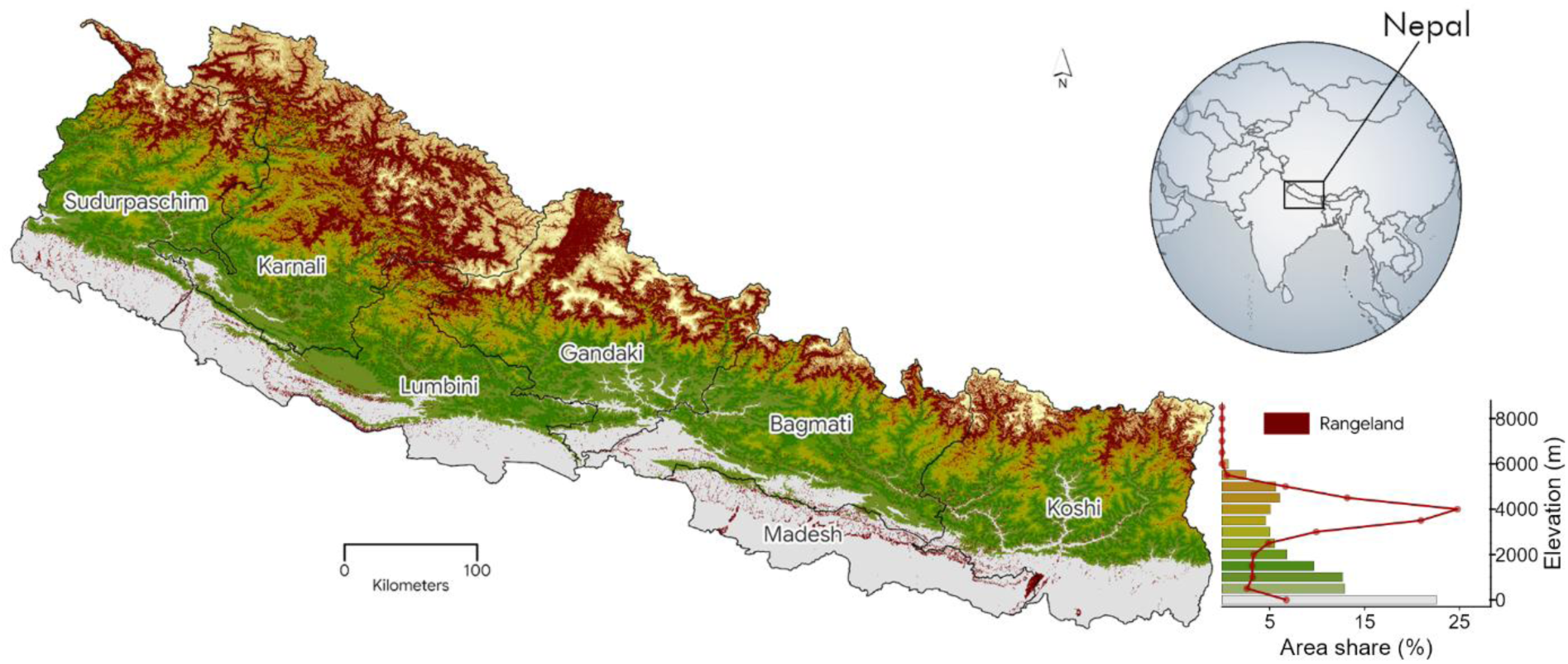
Spatial distribution of rangelands in Nepal. The bar plot shows the distribution of total land area and rangeland area across elevation bands.

### 2.2. Normalized Difference Vegetation Index (NDVI)

Normalized Difference Vegetation Index (NDVI) from 2000 to 2025 was derived from the Terra (MOD13Q1) Moderate Resolution Imaging Spectroradiometer (MODIS) Vegetation Indices Version 6.1 products (Didan 2021). These 16-day composite products with spatial resolution of 250 m are produced by compositing maximum NDVI values from multiple observations while minimizing the effects of cloud contamination and atmospheric aerosols (Didan et al. 2015). We further removed cloud-contaminated, snow-covered and low-quality pixels using the MODIS quality assurance layer prior to analysis. Additionally, pixels with NDVI values less than 0.1 were removed to exclude non-vegetative features, such as waterbodies, rock formations, soil, and developed areas.

For each calendar year, annual maximum NDVI (hereafter, NDVImax) was extracted for each pixel as the highest quality-filtered composite value within that year. NDVImax is a robust indicator of peak growing-season canopy greenness that minimizes the influence of soil background reflectance, residual cloud artefacts, and inter-site phenological offsets (Eckert et al. 2015).

### 2.3. Precipitation and Temperature

Daily precipitation and temperature data were obtained from ERA5-Land daily aggregated data at 11,132 m spatial resolution through Google Earth Engine (GEE) (Hersbach et al. 2020). Daily data were aggregated to monthly precipitation and temperature for each pixel.

### 2.4. RESTREND analysis

Residual Trend Analysis (hereafter, RESTREND) was used to identify NDVI trends not explained by interannual variation in precipitation and/or temperature, and a residual component associated with factors not captured by the modeled climatic predictors (Evans & Geerken 2004). If NDVI residuals show no systematic temporal trend, the observed change in vegetation productivity can be attributed primarily to climate variability. A significant negative residual trend indicates a decline in rangeland productivity (decline in vegetation greenness) beyond that expected from climate alone, consistent with potential non-climatic degradation. A significantly positive residual trend indicates productivity improvement (increase in vegetation greenness) above the climate-predicted expectation and may potentially reflect land restoration, reduced grazing pressure, shrub encroachment, or management change or other non-climatic processes (Evans & Geerken 2004; Li et al. 2012; He et al. 2015; Burrell et al. 2017). RESTREND workflow included three major steps: i) identifying optimal climate windows by establishing the climate-vegetation relationship, ii) isolating the climate-predicted vegetation change, iii) isolating residual vegetation change not explained by the modeled climate variables.

A key challenge in RESTREND analysis is that the climatic window most strongly associated with peak vegetation productivity can vary across pixels because of differences in phenology, elevation, and land cover type (Burrell et al. 2017). To identify the precipitation accumulation period and temperature averaging window that strongly predict NDVImax at each pixel, we first determined the typical timing of peak greenness across Nepal’s rangelands using TIMESAT software version 3.3 (Jönsson & Eklundh 2004). The day-of-year at which NDVImax occurred was mapped for 2013 (the approximate midpoint of the 2000–2025 study period) across all rangeland pixels. The resulting day-of-year image was spatially averaged across all retained rangeland pixels to identify a reference month. This analysis identified August as the month of peak productivity across Nepal’s rangelands (Supplementary Figure S1), consistent with the late-monsoon phenological pattern documented in Himalayan alpine and sub-alpine grassland systems.

Because the timing of NDVImax varies among years and along elevation gradients, climate windows ending two to three months before and after August were also evaluated. Starting from January, cumulative precipitation windows and average temperature windows of varying durations were systematically generated for each reference end month (June, July, August, September, October, and November), together with single-month windows to capture short-term lagged responses. This procedure yielded 56 candidate climate windows for each pixel (Supplementary Table S1).

Following Burrell et al. (2017, 2019), two complementary RESTREND models were fitted across all rangeland pixels, representing two hypotheses about the primary climatic limitation on vegetation productivity. The first was Vegetation-Precipitation Relationship (VPR) model, in which NDVImax was regressed against the optimal accumulated precipitation only. This model is appropriate for arid and semi-arid pixels where moisture availability is the dominant control on vegetation productivity and temperature plays a secondary role (Burrell et al. 2017). The second was Vegetation-Climate Relationship (VCR) model, in which NDVImax was regressed against optimal accumulated precipitation, optimal mean temperature, and their interaction term at each pixel. This model is appropriate for cold, high-altitude environments where low temperature can constrain photosynthesis even when moisture is adequate, and for pixels where the combined thermal-moisture environment best predicts vegetation productivity (Burrell et al. 2019; Gao et al. 2022).

For the VPR model, the regression at each pixel was:

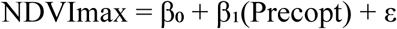

For the VCR model, the regression was:

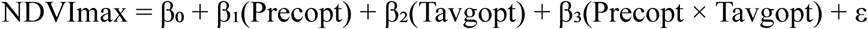

where Precopt and Tavgopt are the pixel-specific optimal precipitation and temperature time series respectively; β₀ is the intercept; β₁–β₃ are regression coefficients; and ε is the model error term.

For VPR model, all 56 candidate precipitation windows were iteratively correlated with the NDVImax time series using Pearson’s correlation coefficient. The precipitation window with the highest significant positive correlation with NDVImax was selected as the optimal precipitation predictor for that pixel (r > 0, p < 0.05), enforcing the ecological assumption that increased moisture supports rangeland productivity in water-limited environments.

For the VCR model, paired precipitation and temperature windows of identical duration were evaluated against NDVImax using multiple correlation analysis. The paired window with the highest significant positive correlation with NDVImax was selected as the optimal joint climate predictor for that pixel (r > 0, p < 0.05).

After the optimal climate predictor was identified for each pixel, only pixels with statistically significant climate–NDVI relationships (p < 0.05) were retained from each model. For retained pixels, both VPR and VCR models were fitted using the ridge regression function in GEE, with the regularization parameter set to 0. This setting is equivalent to ordinary least squares regression (García et al. 2016; McDonald 2009; Yildirim & Revan Özkale 2019). The fitted models were then used to predict NDVImax.

For each retained pixel and each year, NDVI residuals were calculated as observed NDVImax minus predicted NDVImax. To characterize the direction and magnitude of observed NDVImax trends and residual NDVI trends over 2000–2025, we applied the non-parametric Mann-Kendall trend test with Sen’s slope estimator (Mann 1945; Kendall 1975; Sen 1968) using the pymannkendall package in Python. The Mann-Kendall test is widely used in satellite-based vegetation trend analysis because it does not assume normality, is robust to outliers, and is suitable for detecting monotonic trends in environmental time series (Eckert et al. 2015). Sen’s slope provides a robust non-parametric estimate of the rate of change, expressed as NDVI or NDVI residual change per year.

To produce a single spatially continuous trend maps (predicted NDVImax and NDVI residuals), results from the VPR and VCR models were combined into a final mosaic. For pixels where both models met the significance (p < 0.05), the trend from the model with the higher correlation coefficient between optimum climate and predicted NDVImax was selected. This model was considered to provide a stronger explanation of climate-related NDVI variance and, therefore, more reliable results. For pixels where only one model met the thresholds, the trend from that model was used. Pixels where neither model met the thresholds were classified as indeterminate and excluded from change attribution.

Pixels were classified into two final trend categories based on the sign and significance of Sen’s slope: significant positive trend, defined as p < 0.05 with a positive Sen’s slope; and significant negative trend, defined as p < 0.05 with a negative Sen’s slope.

In addition to isolating climatic-predicted and residual vegetation trends using RESTREND, we evaluated observed vegetation trends in annual NDVImax over the 26 year-study period using Mann-Kendall trend test with Sen’s slope estimator. For each pixel, the sign and statistical significance of Sen’s slope were used to classify observed NDVImax trends into two categories: significant positive trend and significant negative trend (p < 0.05). Pixels with non-significant trends were classified as no significant change.

### 2.5. Livestock data

Annual livestock distribution data for cattle, goats, and sheep at 1-km resolution for 2000-2022 were obtained from Global Pasture Watch (Parente et al. 2025). Each pixel value represents the number of animals calibrated to country-level FAOSTAT livestock statistics using dasymetric spatial disaggregation. This dataset provides spatially consistent and temporally resolved gridded livestock data currently available for Nepal.

Livestock counts for cattle, goats, and sheep were converted to standardized animal units (AU) using species-specific animal unit equivalency (AUE) coefficients. We used 1.0 AU for cattle, 0.1 AUs for sheep and goats (Redfearn & Bidwell 2017). Total AU per pixel per year was calculated as: Total AU = (Ncattle × 1.0) + (Ngoats × 0.1) + (Nsheep × 0.1).

Pixel-wise temporal trends in total AU over 2000–2022 were then characterized using the Mann-Kendall trend test with Sen’s slope estimator. Pixels were classified as having a significant increase, significant decrease, or no significant trend in animal units.

### 2.6. Spatial comparison of rangeland and livestock trends

To assess the spatial correspondence between areas of rangeland change and livestock pressure, we conducted the analysis at municipality scale by spatially aggregating both datasets at the municipality level. For each municipality, we calculated the area or proportion of rangeland pixels showing significant residual positive and negative NDVI trends and the area or proportion of livestock pixels showing significant AU increase or decrease. These values were compared to assess broader administrative-level patterns between vegetation change and livestock pressure. All data processing was conducted using the Google Earth Engine (GEE) Python API (Gorelick et al. 2017).

## 3. Results

### 3.1. Extent and distribution of Nepal’s rangelands

Stable rangelands, defined from annual LULC maps using 25th-percentile temporal agreement, covered approximately 16.4% (2,415,270 ha) of Nepal’s total land area. Most rangelands occurred outside protected areas spanning at an elevation range of 2,000 to over 7,000 m above sea level, representing 85.34% (2,058,793 ha) of the stable rangeland area.

Among Nepal’s seven provinces, Karnali Province contained the largest share, with 893,667 ha (43.4%) of rangelands, reflecting Karnali’s concentration of vast trans-Himalayan and high-alpine pastoral landscapes in districts such as Dolpa, Humla, and Mugu. Gandaki Province had the second largest share (463,326 ha; 22.5%), followed by Sudurpashchim (262,782 ha; 12.8%), Koshi (247,244 ha; 12.0%), Bagmati (154,162 ha; 7.5%), and Lumbini (37,613 ha; 1.8%) (Figure 2a). Madhesh Province contained no rangeland above 2,000m, consistent with its lowland Tarai geography.

**Figure 2.**
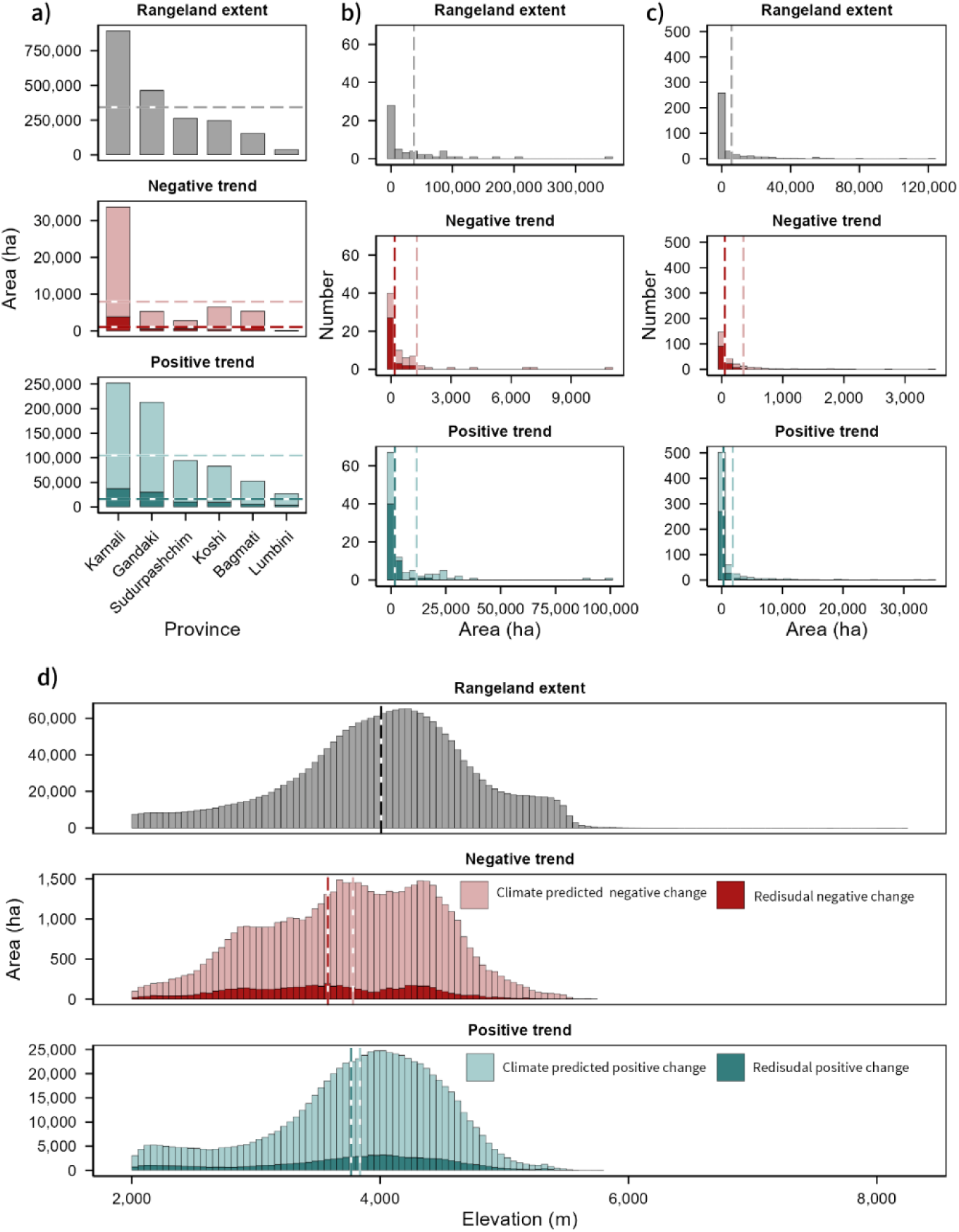
Distribution of rangeland extent, climate-predicted NDVI trends, and residual NDVI trends across administrative and elevational units: (a) province, (b) district, (c) municipality, and (d) elevation band.

Rangelands were concentrated mainly in the subalpine and alpine zones, peaking at approximately 4,000–4,500 m, with a mean elevation of approximately 4,200 m (Figure 2d). Rangelands below 3,000 m represented a relatively small fraction of the total, while rangelands above 5,500 m were sparsely distributed. Out of Nepal’s 77 districts, 52 districts had rangelands above 2000m (Figure 2b). Of the 753 local government units, 285 municipalities had rangelands above 2000m (Figure 2c).

### 3.2. Spatial pattern of rangeland change

Mann-Kendall trend analysis of annual maximum NDVI across Nepal’s 2,058,793 ha of non-protected rangelands revealed that the majority of rangelands (80.7%; 1,660,811 ha) showed no statistically significant vegetation change over the last 26 years (Figure 3a). Significant NDVImax trends were detected across 397,983 ha (19.3%), of which 383,281 ha (96.3%) showed positive trends or increase in vegetation greenness, whereas 14,702 ha (3.7%) showed negative trends or decline in vegetation greenness. Expressed relative to total rangeland area, positive annual maximum NDVI trends covered 18.6%, while negative trends covered 0.7%.

**Figure 3.**
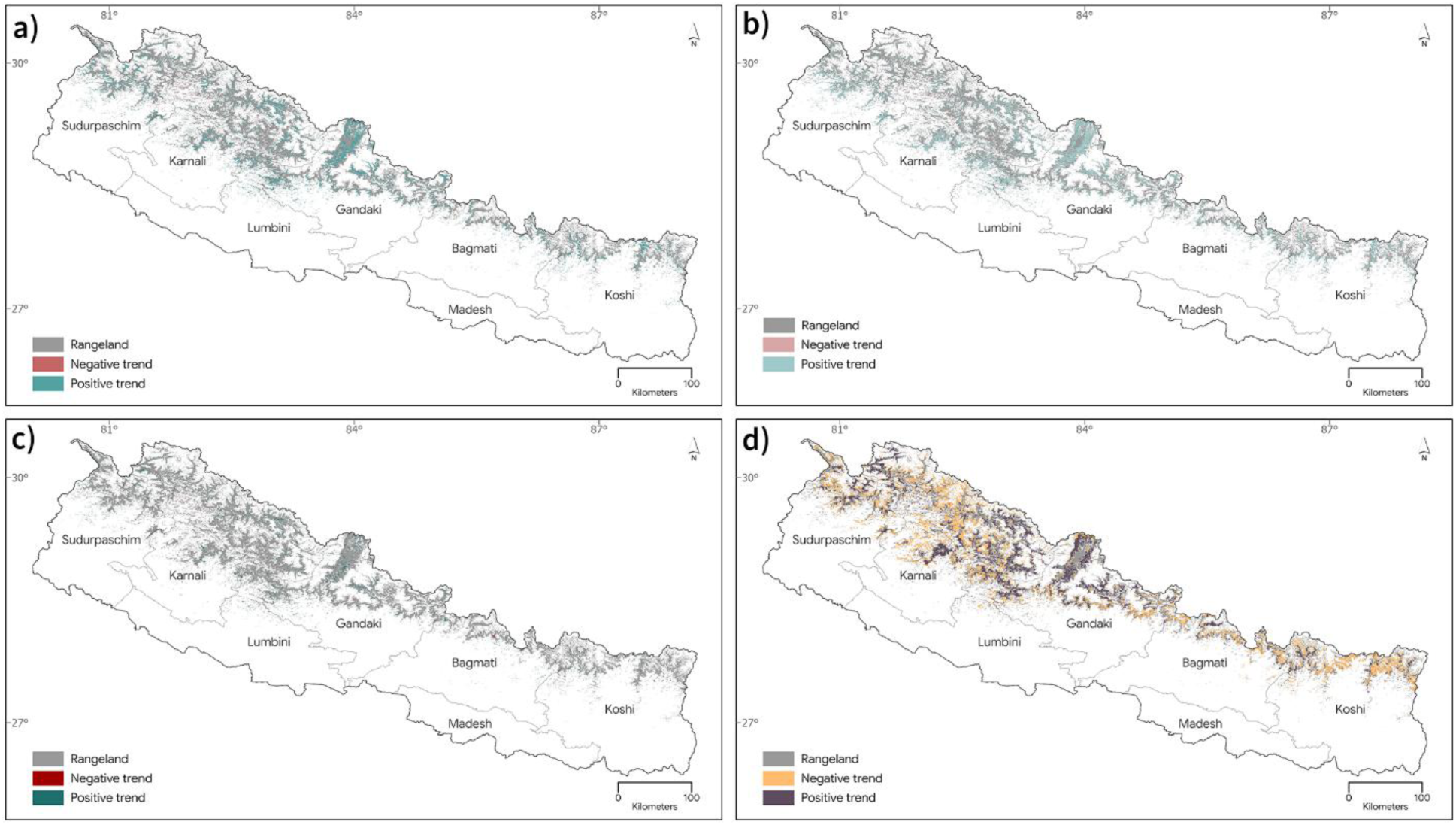
Spatial patterns of rangeland vegetation change and livestock trends: (a) observed annual maximum NDVI trends, (b) climate-predicted NDVI trends, (c) residual NDVI trends, and (d) animal unit trends.

At the provincial scale, Lumbini had the highest proportional positive NDVImax (37.8%) followed by Gandaki (28.1%) and Sudurpashchim (18.3%). Karnali (13.8%) had a lower proportional positive trend but one of the largest absolute areas of change because it contained the largest rangeland extent (Table 1). For negative annual maximum NDVI trends, Bagmati had the highest proportional decline in vegetation greenness (1.4%), followed by Karnali (1.1%), whereas Lumbini had the lowest decline in vegetation greenness (0.1%) (Table 1).

**Table 1.**
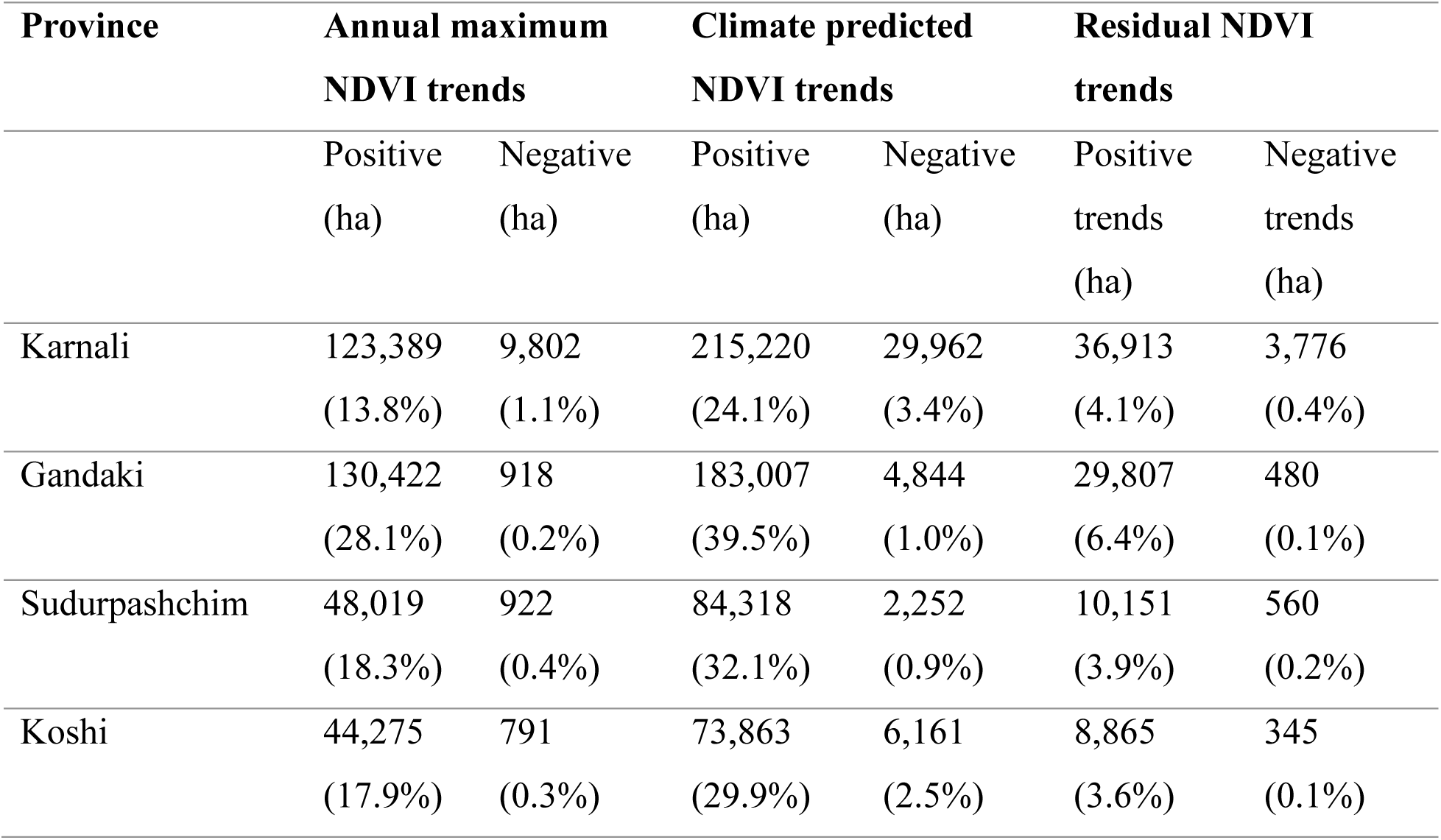
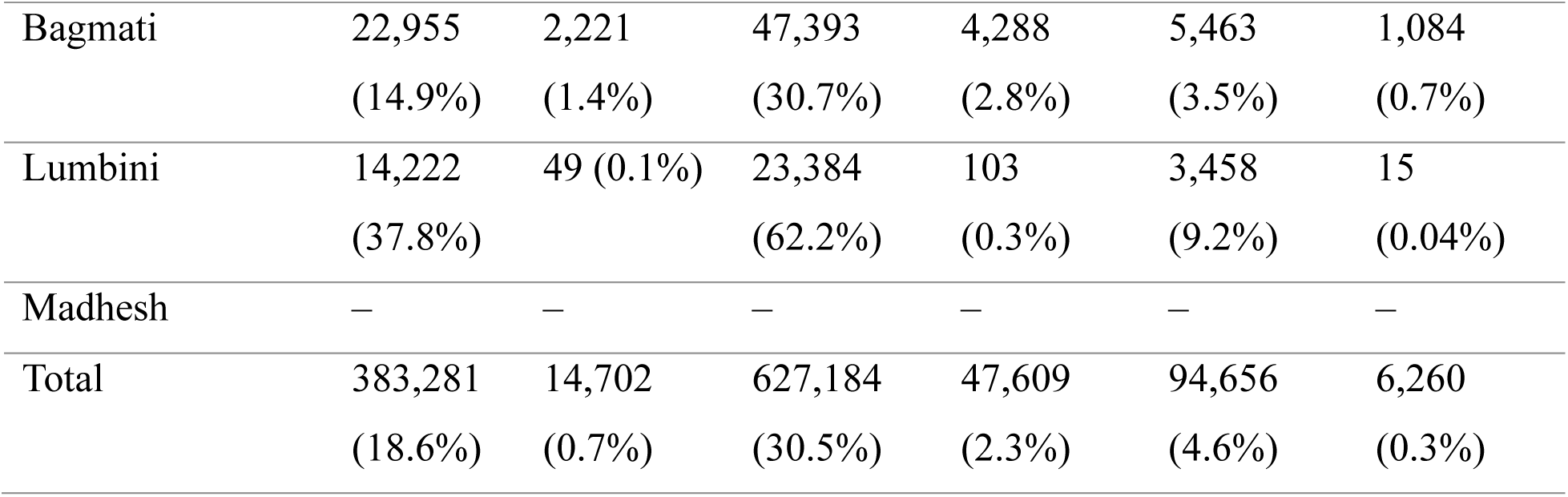
Province-wise area of annual maximum NDVI trends, climate-predicted NDVI trends, and residual NDVI trends.

RESTREND analysis showed that the climate-predicted NDVI component was spatially more extensive than the observed annual maximum NDVI trend. Climate-predicted positive trends covered 627,184 ha (30.5% of total rangeland area), whereas residual positive trends covered 94,656 ha (4.6%). Among the 674,793 ha of significant climate-predicted NDVI change, 627,184 ha (92.9%) showed positive trends and 47,609 ha (7.1%) showed negative trends (Figure 3).

At the provincial scale, Lumbini had the highest proportional climate-predicted positive trend (62.2%), followed by Gandaki (39.5%), Sudurpashchim (32.1%), Bagmati (30.7%), Koshi (29.9%), and Karnali (24.1%) (Table 1, Figure 3c). In absolute area, however, Karnali, followed by Gandaki and Sudurpashchim, contributed the largest areas of climate predicted positive trend.

Climate-predicted negative trends covered 47,609 ha (2.3%) of total rangeland area, compared with 6,260 ha (0.3%) under residual negative trends. This indicates that negative NDVI trends were more spatially extensive in the climate-predicted component than in the residual component. For residual positive trends, Lumbini recorded the highest proportional area (9.2%), followed by Gandaki (6.4%), Karnali (4.1%), Sudurpashchim (3.9%), Koshi (3.6%), and Bagmati (3.5%). For residual negative trends, Bagmati recorded the highest proportional area (0.7%), followed by Karnali (0.4%) (Figure 3b).

At the district scale, 36 districts showed climate-predicted negative trends while 52 districts showed climate-predicted positive trends, mostly occurred in high mountain districts. Climate-predicted positive change was greatest in Dolpa (99,354 ha; 28.0%), Mustang (88,634 ha; 50.8%), Bajhang (37,132 ha; 35.9%), Humla (30,342 ha; 14.2%), and Manang (29,961 ha; 37.8%). Climate-predicted negative change was largest in Dolpa (10,888 ha; 3.1%), Humla (6,955 ha; 3.3%), Mugu (6,653 ha; 5.1%), Jumla (4,133 ha; 3.8%), and Sankhuwasabha (2,968 ha; 4.6%) (Supplementary Figure S2).

For residual positive trends, the largest areas occurred in Dolpa (15,913 ha; 4.5%), Mustang (14,363 ha; 8.2%), Humla (8,688 ha; 4.1%), Manang (5,133 ha; 6.5%), and Bajhang (5,082 ha; 5%). Residual negative trends were more limited in extent, with the largest areas in Humla (1,288 ha, 0.6%), Dolpa (1,040 ha, 0.3%), Mugu (860 ha, 0.7%), Sindhupalchok (661 ha, 1.7%), and Jumla (388 ha, 0.4%) (Supplementary Figure S3).

Among Nepal’s 753 municipalities, 285 contained rangelands with climate-predicted positive trends while 121 municipalities had rangelands with climate-predicted negative trend. Residual positive trends were observed in 281 municipalities, whereas residual negative trends occurred in 121 municipalities.

Across elevation band, RESTREND results showed that both climate-predicted and residual trends were concentrated mainly between 3,000 and 5,000 m. Climate-predicted positive change was largest in the 4,000–5,000 m band (283,884 ha, 40.5%), followed by the 3,000–4,000 m band (278,793 ha, 39.8%). Climate-predicted negative change was also greatest in the 3,000–4,000 m band (20,699 ha, 3.0%) and the 4,000–5,000 m band (18,466 ha, 2.6%). Residual positive change followed a similar pattern, with the largest areas in the 4,000–5,000 m band (39,816 ha, 38.5%) and the 3,000–4,000 m band (39,434 ha, 38.2%). Residual negative change was also concentrated mainly in the 3,000–4,000 m band (2,823 ha, 2.7%) and the 4,000–5,000 m band (1,919 ha, 1.9%).

### 3.3. Livestock trends and correspondence with rangeland change

Trends in total livestock animal units (AU) over 2000–2022 showed significant spatial variation across Nepal’s rangelands. Significant increasing AU trends were overlapped with 523,182 ha (25.4%) of total rangeland area, while significant decreasing AU trends overlapped with 395,376 ha (19.2%). Thus, the area experiencing increasing livestock pressure was larger than the area experiencing declining livestock pressure. However, a substantial proportion of rangelands showed no significant AU trend or remained outside the classified AU trend categories, indicating spatial heterogeneity in livestock pressure across Nepal’s high-altitude rangelands.

Comparison of municipality-averaged AU trends with RESTREND residual trends suggested that livestock pressure alone did not explain the observed residual change caused by non-climatic drivers in Nepal’s high-altitude rangelands (Figure 3d). Residual positive NDVI trends covered 94,656 ha (4.6%), whereas residual negative trends covered only 6,260 ha (0.3%). If livestock pressure were the dominant driver of residual vegetation trends, municipalities with increasing AU would be expected to show stronger residual NDVI decline, while municipalities with decreasing AU would be expected to show residual NDVI increase. However, no clear linear relationship between AU trends and residual NDVI trends suggesting that municipalities were widely distributed across all four comparisons, with weak fitted relationships (Figure 4). This suggests that livestock pressure may contribute to non-climatic rangeland change in some locations, but it is not the sole or dominant non-climatic driver of rangeland change across Nepal.

**Figure 4.**
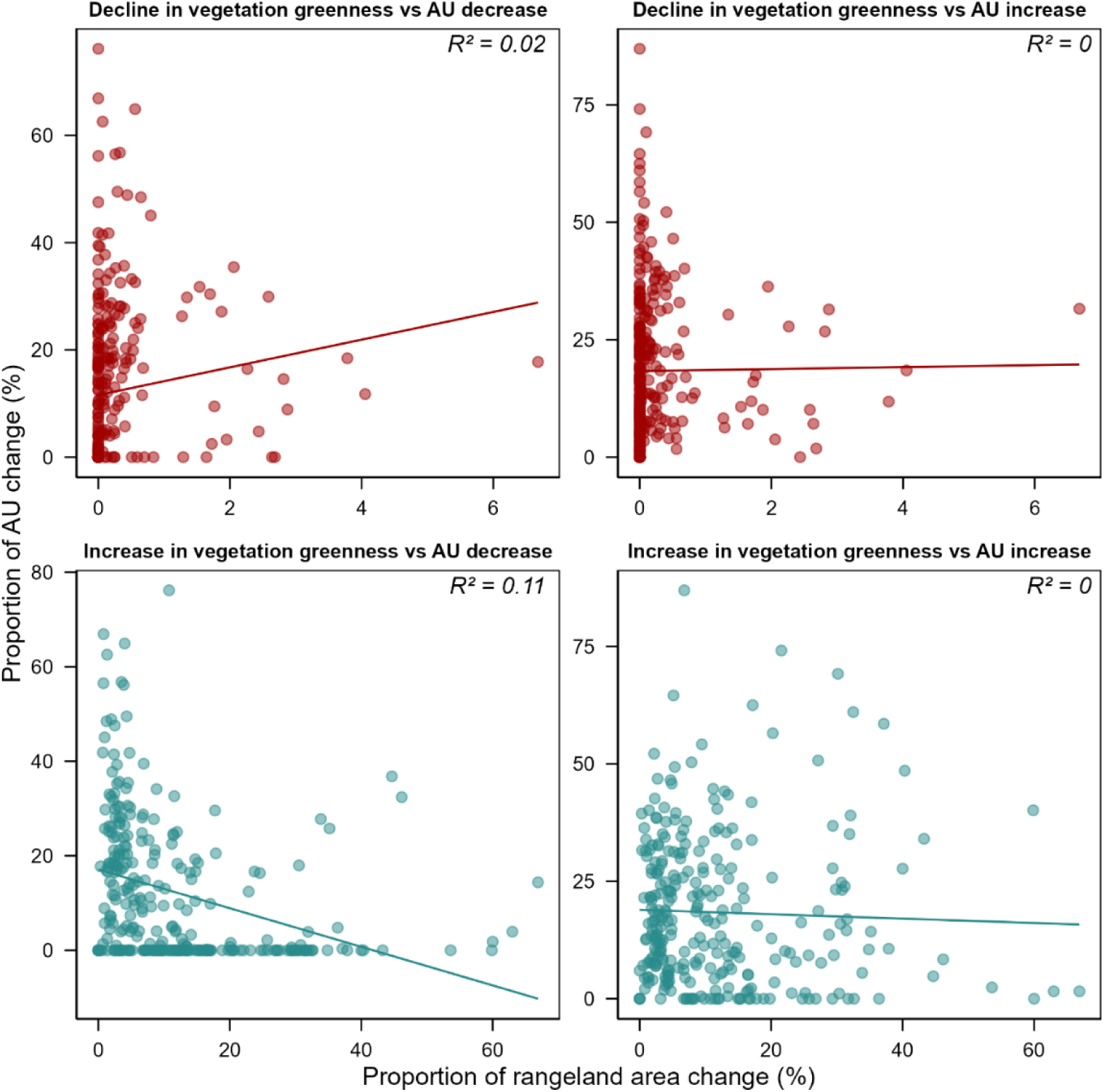
Relationship between rangeland residual NDVI trends and livestock animal unit trends at the municipality scale.

## 4. Discussion

This study provides a multi-scale RESTREND-based assessment of vegetation change across Nepal’s non-protected rangeland ecosystems, examining the effect of climate-predicted and residual NDVI trends at municipality, district, province, and elevation levels. The results show that most rangelands remained stable over the study period, with positive annual maximum NDVI trends substantially more extensive than negative trends. Climate-predicted NDVI trends were more spatially extensive than residual NDVI trends, suggesting that climate-related variability explained a larger share of rangeland vegetation dynamics than residual non-climatic processes at the national scale. Residual negative trends were limited in extent and spatially concentrated in selected high mountain and peri-urban areas. Municipality-level livestock trends did not explain the residual NDVI signal, indicating that non-climatic vegetation change likely reflects multiple interacting drivers, including grazing dynamics, land-use change, shrub expansion, invasive species, and limitations in gridded livestock data.

The finding that 80.7% of Nepal’s non-protected rangelands showed no significant annual maximum NDVI trend, and that significant change is dominated by positive trends (96.3%) rather than negative trends (3.7%), is consistent with broader vegetation trends reported across Nepal and the Hindu Kush Himalayan region. Previous studies reported increasing NDVI trends in Nepal from 1982-2015 (Baniya et al. 2020) and 2000-2019 (Nila et al. 2025) and about 80% of Nepal’s vegetation area showed increasing NDVI trends from 2003 to 2022 (Shrestha et al. 2024). Localized study based on MODIS NDVI product also reported increasing NDVI trends in central Nepal (Regmi et al. 2020) and eastern Nepal (Jian et al. 2025). These studies identified climate, particularly precipitation and warming, together with the success of community forestry program, as contributors to increasing vegetation productivity (Baniya et al. 2020; Shrestha et al. 2024; Jian et al. 2025). In Nepal, NDVI has been reported to correlate more strongly with temperature than with precipitation (Baniya et al. 2020; Shrestha et al. 2024). Higher elevation where rangelands are the dominated vegetation type in Nepal, it is the temperature rather than precipitation that enhance vegetation growth (Nila et al. 2025; Shrestha et al. 2024). In higher-elevation areas, where rangelands dominate, temperature rather than precipitation may be the stronger control on vegetation growth (Nila et al. 2025; Shrestha et al. 2024). Katel et al. (2026) and Shrestha et al. (2012) also reported a lengthening growing season in alpine and subalpine vegetation in Nepal and the Himalaya respectively. Warming-driven extension of the growing season therefore provides a plausible mechanism for the widespread positive NDVI signal observed across Nepal’s rangelands in this study.

The RESTREND framework extends simple NDVI trend analysis by separating observed vegetation variation into a climate-predicted component and a residual component not explained by the modeled climate variables (Evans & Geerken 2004; Li et al. 2012). In this study, climate-predicted trends were more extensive than residual trends for both positive and negative NDVI changes. This pattern suggests that climate-related variability was the dominant broad-scale signal in Nepal’s rangelands, whereas trends predicted by non-climatic drivers were more localized. However, residual positive trends should not be interpreted automatically as improved rangeland condition. They may reflect genuine recovery from historical grazing pressure, but they may also indicate shrub encroachment, changes in species composition, land-use change, or pastoral abandonment or management interventions that increase NDVI without necessarily improving forage quality or pastoral value.

This finding aligns closely with RESTREND-based studies across dryland and mountain systems globally. Lehnert et al. (2016) found that climate variability, including precipitation declines in the central and western Tibetan Plateau and warming-induced moisture stress, was the primary driver of large-scale vegetation cover change, while increasing livestock numbers intensified negative trends locally but were not the dominant regional driver. Wu et al. (2021), using generalized additive mixed modelling (GAMM) and RESTREND on the Qinghai-Tibetan Plateau, found that combined climatic variables, including temperature, precipitation, and radiation, explained 66.2% of NDVI variance at the pixel scale, whereas human influence affected 15.5–14.3% of grassland pixels. These findings are consistent with our result that climate-predicted NDVI trends were more extensive than residual trends in Nepal’s rangelands.

RESTREND studies in other dryland systems also support the importance of climate as a major driver of vegetation dynamics. Gedefaw et al. (2021) applied TSS-RESTREND to New Mexican rangelands and found that 17.6% experienced productivity decline, with breakpoints coinciding with drought events indexed by the Palmer Drought Severity Index. Chen et al. (2023) showed that climate change accounted for 48.3% of vegetation change in Chinese drylands using TSS-RESTREND. Zhuge et al. (2019) found that climate change was the main driver of land degradation in northern China over three decades, while human impacts were more prominent at smaller spatial and temporal scales. These studies support our interpretation that climate-predicted factors can dominate rangeland vegetation change at landscape to national scales, while grazing and other non-climatic processes may be more important locally. This local-scale complexity is consistent with Paudel and Andersen (2010), who reported that grazing-induced change and annual precipitation variability can both shape rangeland vegetation dynamics. In our study, municipality-level analysis identified locations such as Adanchuli, Tanjakot, and Soru where combined climate-predicted and residual negative trends affected 20–26% of rangeland area, indicating localized areas where interacting drivers may require closer field assessment.

We did not find a clear relationship between livestock trends and residual NDVI trends at the municipality scale. The weak relationship may reflect several factors. First, shrub expansion or invasion by species with higher photosynthetic activity than native grasses can increase NDVI even where forage value declines (Cai et al. 2015; Reeves et al. 2024). Shrub encroachment and invasive species expansion have been reported in Nepal’s high-elevation rangelands (Sharma et al. 2014; MoITEF 2026), and conversion of alpine meadows into shrublands has been reported elsewhere in the Himalaya (Brandt et al. 2013). This may explain why some areas with increasing livestock pressure nevertheless showed stable or positive residual NDVI trends. Positive RESTREND residuals in such settings may not indicate improved rangeland condition for livestock production. Instead, they may reflect compositional shifts toward shrub-dominated cover that increase NDVI but reduce forage quality and pastoral value (Brandt et al. 2013; Gxasheka et al. 2023). Distinguishing productive grassland recovery from shrub-driven NDVI increase is a limitation of NDVI-based RESTREND analysis and requires field validation, spectral unmixing, or higher-resolution vegetation composition data. Second, rangeland recovery may not occur even where livestock numbers decline if grazing pressure remains above carrying capacity or if vegetation degradation has crossed ecological thresholds (Zhang et al. 2018).

In this study, the combined area of significant residual trends and climate-predicted trends exceeded the area of significant observed annual maximum NDVI trends. This does not indicate an inconsistency in the analysis. Annual maximum NDVI trends, climate-predicted trends, and residual trends represent different statistical components of the NDVI signal. Annual maximum NDVI trends describe the net observed vegetation response. RESTREND separates this signal into a climate-predicted component and a residual component. These components can act in the same or opposite directions at the same pixel. For example, a positive climate-predicted trend may be offset by a negative residual trend, producing a weak or non-significant observed NDVImax trend. Conversely, a residual trend may be significant even when the observed annual maximum NDVI trend is not significant. Therefore, the areas of climate-predicted and residual trends should not be added and compared directly with the area of observed NDVImax trends. They should instead be interpreted as diagnostic components that help identify where climate-related and residual processes contribute to vegetation change (Evans & Geerken 2004; Abel et al. 2018; Burrell et al. 2019; Li et al. 2021).

Several methodological considerations affect interpretation of these findings. RESTREND partitions NDVI trends into climate-predicted and residual components, but this partitioning depends on the strength and quality of the climate-NDVI regression at each pixel. Pixels with weak or non-significant climate-NDVI relationships were classified as indeterminate and excluded from attribution. This reduces the risk of spurious attribution but also limits the spatial coverage of attributed results. The predominance of the VCR model across high-altitude pixels reflects the importance of temperature alongside precipitation in explaining NDVImax variation, consistent with Burrell et al. (2019), who showed that including temperature reduces the fraction of vegetation change incorrectly attributed to neither climate nor land use. However, RESTREND cannot distinguish among different categories of non-climatic change. Shrub encroachment, recovery from overgrazing, land-use conversion, and management interventions can all produce residual positive NDVI trends and cannot be separated without additional field, spectral, or land-cover change data (Chen et al. 2023).

The use of gridded 1 km livestock data (Parente et al. 2025) introduces spatial and temporal uncertainties that limit the precision of livestock attribution. Gridded livestock models are derived through dasymetric disaggregation of census data using environmental covariates and may not capture the seasonal concentration of livestock on specific pastures during transhumance, a pattern that is ecologically important in Nepal’s high-altitude pastoral systems (Luxom et al. 2022; Basnet et al. 2024). The spatial comparison of RESTREND trends with AU trends should therefore be interpreted as evidence of broad-scale patterns in grazing-pressure dynamics rather than definitive causal attribution of NDVI change to livestock. Field validation of RESTREND-classified pixels, particularly in areas with residual negative trends such as Sindhupalchok and Rasuwa and areas with strong residual positive trends such as Mustang and Manang, is needed to evaluate the relative influence of climate-related and non-climatic drivers at the landscape scale.

The overall pattern of residual positive NDVI trends across Nepal’s high-altitude rangelands is encouraging, but its ecological meaning requires caution. Where residual positive trends reflect recovery of productive rangelands from historical overgrazing, the results suggest that reduced stocking pressure can support passive rangeland recovery and that policies promoting appropriate grazing intensity may be beneficial. Where residual positive trends reflect shrub encroachment driven by pastoral abandonment, however, the NDVI signal may be misleading. Replacement of open alpine grasslands by unpalatable shrubs such as *Berberis* may reduce forage value and pastoral benefits even if NDVI increases (Sharma et al. 2014; Brandt et al. 2013). In such areas, active management interventions, including controlled burning, targeted grazing, and weed management, may be required to maintain semi-natural grassland character. The RESTREND-derived classifications presented here provide a spatially explicit evidence base for prioritizing rangeland interventions and for monitoring rangeland vegetation condition in future assessments.

## Author contributions

Conceptualization: UBS; Data curation and Formal analysis: SJ; Methodology: SJ, UBS, Supervision: UBS; Visualization: UBS, SJ; Writing and editing: UBS, SJ.

## Acknowledgements

We sincerely acknowledge the support provided by National Geographic Society (NGS-62058R-9).

