## Supplementary Figure for "Climatic and non-climatic drivers of rangeland vegetation change in Nepal"

**Supplementary Materials**

**Determining the peak of greenness across Nepal’s rangelands**

We first determined the typical timing of peak greenness across Nepal's rangelands using TIMESAT software version 3.3 (Jönsson & Eklundh 2004). The day-of-year at which NDVImax occurred was mapped for 2013 (the approximate midpoint of the 2000–2025 study period) across all rangeland pixels. The resulting day-of-year image was spatially averaged across all retained rangeland pixels to identify a reference month. This analysis identified August as the month of peak productivity across Nepal’s rangelands.

Figure S1. Spatially and temporally averaged NDVImax values curve from 2000 to 2026.

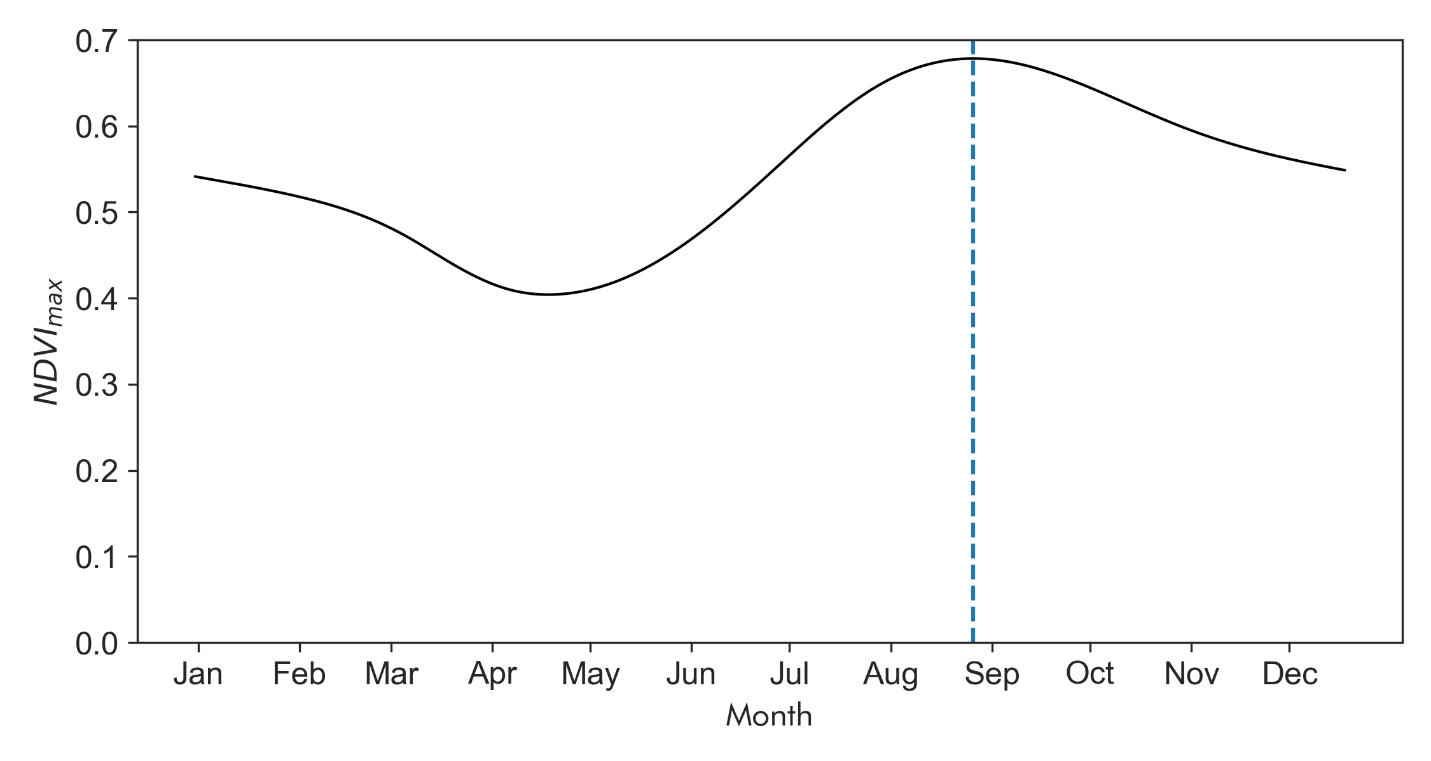

Figure S1. Spatially and temporally averaged annual maximum NDVI values from 2000 to 2025.

Because the timing of NDVImax varies among years and along elevation gradients, climate windows ending two to three months before and after August were also evaluated. Starting from January, cumulative precipitation windows and average temperature windows of varying durations were systematically generated for each reference end month (June, July, August, September, October, and November), together with single-month windows to capture short-term lagged responses. This procedure yielded 56 candidate climate windows for each pixel (Table S1).

| Table S1. Selected 56 climatic windows for VPR and VCR models. | | | | | | |
| --- | --- | --- | --- | --- | --- | --- |
| Single Month Window | Window Ending In | | | | | |
|  | Jun | Jul | Aug | Sep | Oct | Nov |
| Jan | Jan-Jun | Jan-Jul | Jan-Aug | Jan-Sep | Jan-Oct | Jan-Nov |
| Feb | Feb-Jun | Feb-Jul | Feb-Aug | Feb-Sep | Feb-Oct | Feb-Nov |
| Mar | Mar-Jun | Mar-Jul | Mar-Aug | Mar-Sep | Mar-Oct | Mar-Nov |
| Apr | Apr-Jun | Apr-Jul | Apr-Aug | Apr-Sep | Apr-Oct | Apr-Nov |
| May | May-Jun | May-Jul | May-Aug | May-Sep | May-Oct | May-Nov |
| Jun |  | Jun-Jul | Jun-Aug | Jun-Sep | Jun-Oct | Jun-Nov |
| Jul |  |  | Jul-Aug | Jul-Sep | Jul-Oct | Jul-Nov |
| Aug |  |  |  | Aug-Sep | Aug-Oct | Aug-Nov |
| Sep |  |  |  |  | Sep-Oct | Sep-Nov |
| Oct |  |  |  |  |  | Oct-Nov |
| Nov |  |  |  |  |  |  |
| Dec |  |  |  |  |  |  |

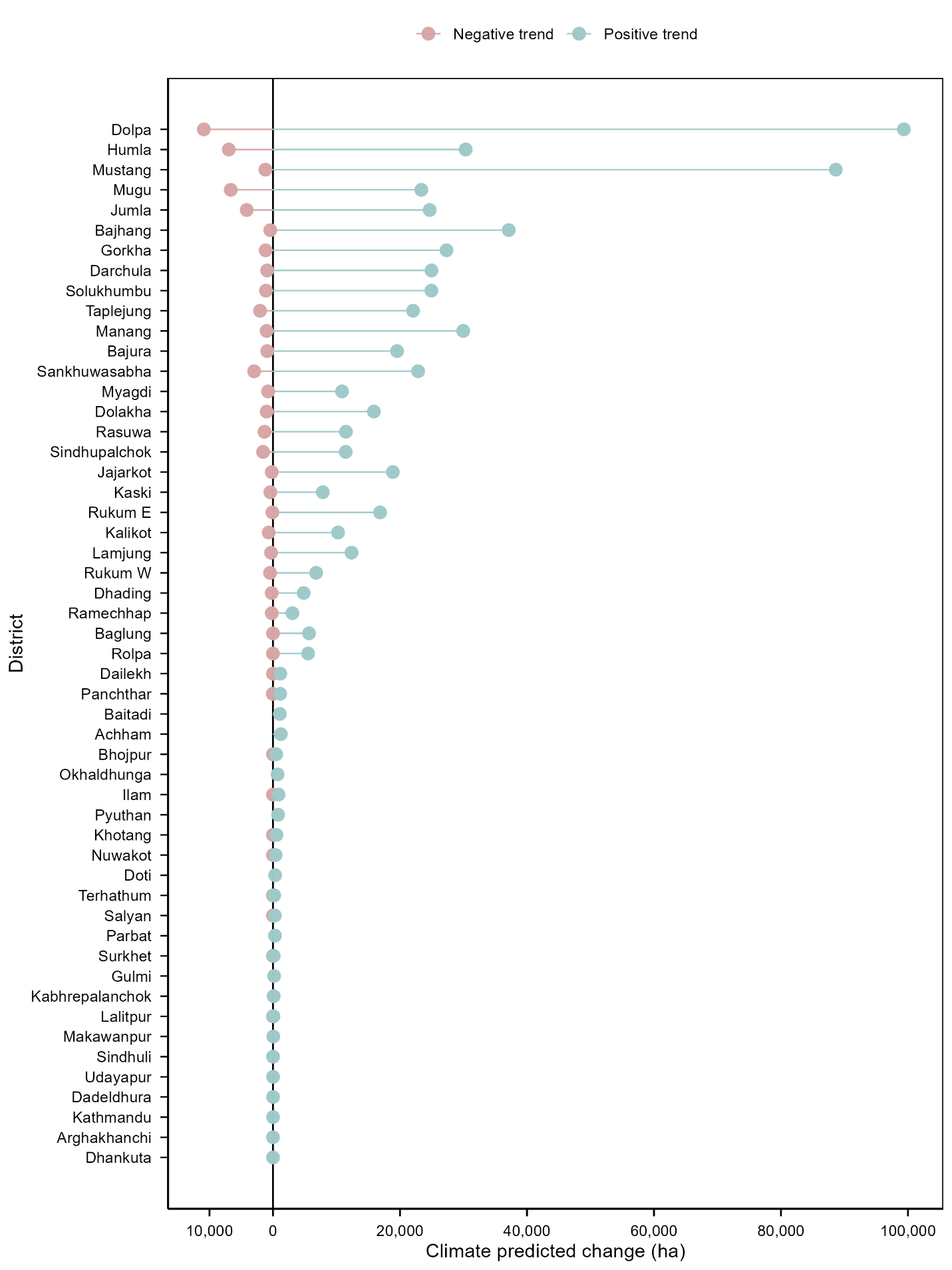

Figure S2. District-wise climate-predicted NDVI trends in rangeland vegetation.

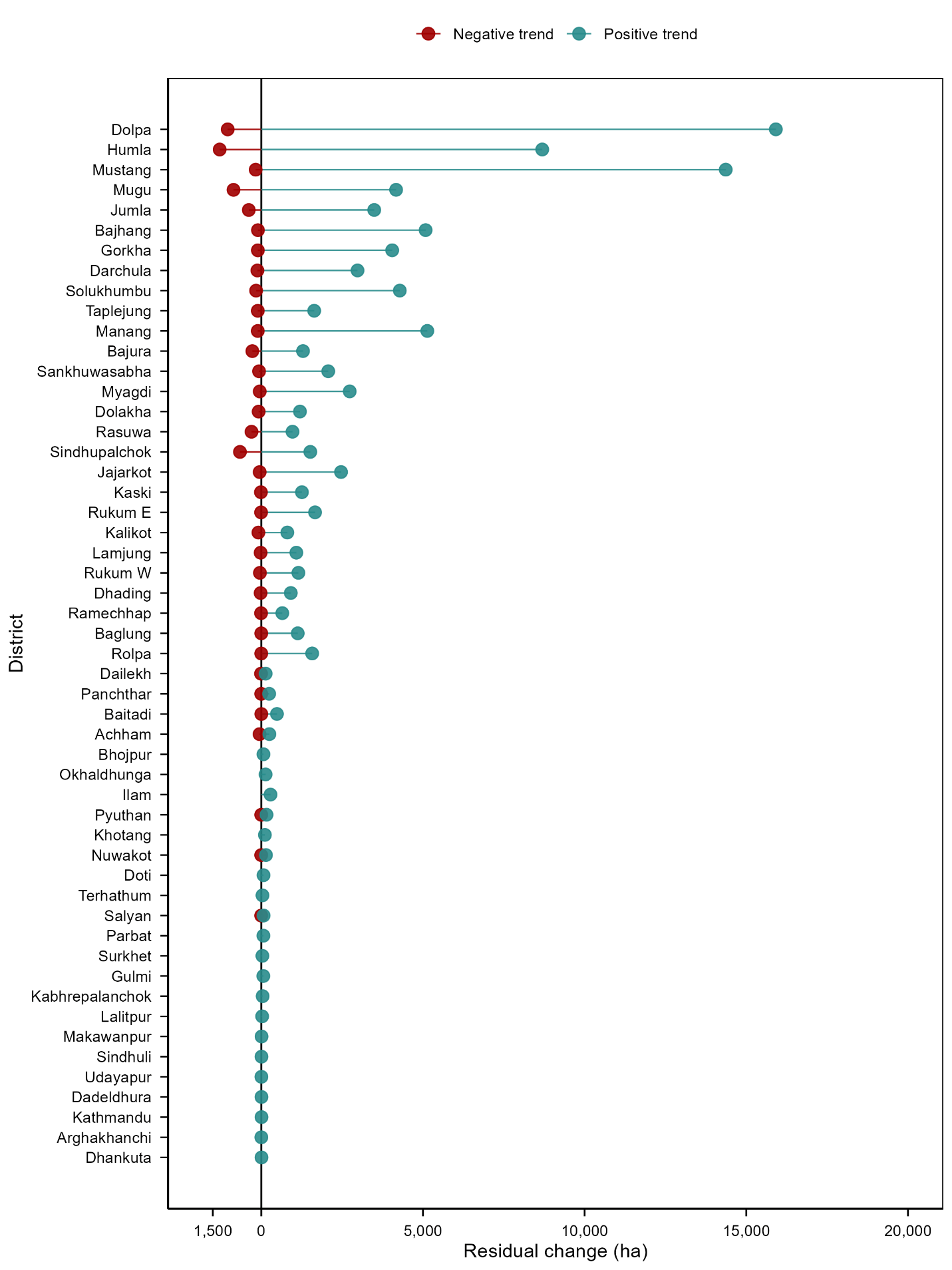

Figure S3. District-wise residual NDVI trends in rangeland vegetation.
